# System Dynamics Modeling of Within-Host Viral Kinetics of Coronavirus (SARS CoV-2)

**DOI:** 10.1101/2020.06.02.129312

**Authors:** Javier Burgos

## Abstract

Mathematical models are being used extensively in the study of SARS-CoV-2 transmission dynamics, becoming an essential tool for decision making concerning disease control. It is now required to understand the mechanisms involved in the interaction between the virus and the immune response effector cells, both innate and adaptive, in order to support lines of research related to the use of drugs, production of protective antibodies and of course, vaccines against SARS-CoV-2. The present study, using a system dynamic approach, hypothesizes over the conditions that characterize the fraction of the population which get infected by SARS-CoV-2 as the asymptomatic patients, the mild symptomatic, acute symptomatic, and also super-spreaders, in terms of innate immune response, the initial virus load, the virus burden with shedding events, and the cytokine levels.

## Introduction

COVID-19 pandemic has underlined the impact of emergent pathogens as a major threat for human health^1,2^. The development of quantitative approaches to advance comprehension of the current outbreak is urgently needed to tackle this severe disease^3^. Coronaviruses (CoV) are a broad family of viruses that can cause a variety of conditions, from the common cold to more serious illnesses, such as the coronavirus that causes Middle East respiratory syndrome (MERS-CoV) and the one that causes the actual respiratory syndrome acute severe (SARS-CoV-2), a new coronavirus that has not been found before in humans^1^.

When an infected person expels virus-laden droplets and someone else inhales them, the novel coronavirus, SARS-CoV-2, enters the nose and throat, and given that the virus binds with a cell-surface receptor called angiotensin-converting enzyme 2 (ACE2), which is distributed throughout the entire body, the virus can disseminate into, potentially, all organs and tissues. As the virus multiplies, an infected person may shed copious amounts of it, especially during the first week or so. Symptoms may be absent at this point. Or the virus’ new victim may develop a fever, dry cough, sore throat, loss of smell and taste, or head and body aches. If the immune system doesn’t beat back SARS-CoV-2 during this initial phase, the virus then marches down the windpipe to attack the lungs, where it can turn deadly. The thinner, distant branches of the lung’s respiratory tree end in tiny air sacs called alveoli, each lined by a single layer of cells that are also rich in ACE2 receptors^4^.

This series of events can be explained taking into account that during an infection, the innate immunity is the first to be triggered (the inflammatory reaction), taking no longer than minutes to hours to be fully activated^5^. This is crucial for the host defense in the first phase of a new infection and plays a central role in the response against SARS-CoV-2, which can explain why most of the infected people will never develop COVID-19 or present mild symptoms (80.9%), the innate immunity is generally able to eliminate pathogens efficiently, but if this initial clearance of SARS-CoV-2 fails, probably due to a high of virus load or virulence of invading pathogens, lymphocytes and adaptative immune mechanisms are activated, increasing antibody-secreting cells (ASCs), follicular helper T cells (TFH cells), activated CD4+ T cells and CD8+ T cells and immunoglobulin M (IgM) and IgG antibodies that bound the COVID-19 causing coronavirus SARS-CoV-2 were detected in blood before symptomatic recovery. Also Accumulating evidence suggests that a subgroup of patients with severe COVID-19 might have a cytokine storm syndrome. However, 13.8% of the infected people develop severe symptoms, 4.4% become critically ill and 2.9% die. These patients usually had chronic diseases, including cardiovascular and cerebrovascular diseases, endocrine system disease, digestive system disease, respiratory system disease, malignant tumors, and nervous system disease^6^.

With the above mentioned, there is an urgent need for understanding the within-host interactions between immune system and SARS-CoV-2, in order to put on the correct framework the results of extensive diagnostic trials all around the world to detect asymptomatic individuals and, of course, to develop novel therapeutics, including antivirals and vaccines^7,8^. Moreover, variables pertaining to the pathogen-host interaction are practically unknown^9^, including the following: dynamic of viral shedding, the role of initial viral load, the dynamic of the viral burden and its relation with the occurrence of cytokine storm, and the relation of these variables with the ongoing of the COVID-19 from asymptomatic towards severe or critically affected. The present study hypothesizes over the conditions that characterize the fraction of the population which get infected by SARS-CoV2, remains asymptomatic or mildly infected, while another fraction being severely symptomatic, the definition of asymptomatic-spreaders and super-spreaders status regarding SARS-CoV-2 infection in humans. Also, shed some light on the dynamics of the Cytokine storm syndrome associated with COVID-19.

## Mathematical model

The present model consists of two components, the first one corresponds to the interaction between two populations, the immune cells and the virus SARS-CoV-2; the second part corresponds to Cytokine release as a response to the antigenic stimulus induced by the interaction between the immune system and SARS-CoV-2 virus.

### Two species interaction

This is a two populations model where *i* and *v* represent, respectively, the immune cells and virus populations at time t= 14 days, which corresponds to the Incubation period (from infection to symptoms). Following Lauer et. al.^10^, the median incubation period is estimated to be 5.1 days (95% CI, 4.5 to 5.8 days), and 97.5% of those who develop symptoms will do so within 11.5 days (CI, 8.2 to 15.6 days) of infection. These estimates imply that, under conservative assumptions, 101 out of every 10 000 cases (99th percentile, 482) will develop symptoms after 14 days (336 hours) of active monitoring or quarantine.

In constructing a model of the interaction of the two populations, the following assumptions are made.

1. In the absence of the immune response, which could be innate immune response (HR) or adaptative immune response, the viral load grows at a rate proportional to the current population; thus 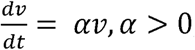, when *i* – 0 and *IIR*=0.
2. In the absence of the virus load the immune cells dies out; thus 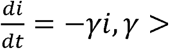, when *v* – 0.
3. The number of encounters between the immune cells and virus is proportional to the product of their populations. Each of such encounters tends to promote the growth of the immune cells, by clonal expansion, and to inhibit the growth of the viral burden. Thus, the growth rate of the immune cells is increased by a term of the form *βvi*, while the growth rate of the viral burden is decreased by a term —*βvi*, and *β* is a positive constant.

As a consequence of these assumptions, it is possible to obtain the set of equations described as follows, starting with the population of viral particles or viral burden (*v*).

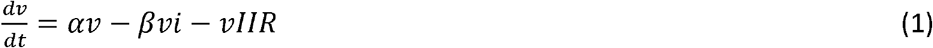

Equation (1) represent the variation in time of the virus population within the host as function of *αv*, being *α* the initial viral load. The last part of the equation (1) means the viral decay by the action of the Innate Immune Response (*HR*); This is crucial for the host defense in the first phase of a new infection, and plays a central role in the response against SARS-CoV-2^9,11,12^, which can explain why most of the infected people will never develop COVID-19 or present mild symptoms (80.9%); Thus, the innate immunity is generally able to eliminate pathogens efficiently.

Moreover, the host innate immune system detects viral infections by using pattern recognition receptors (PRRs) to identify pathogen-associated molecular patterns (PAMPs). At present, the known PRRs mainly include toll-like receptor (TLR), RIG-I-like receptor (RLR), NOD-like receptor (NLR), C-type lectin-like receptors (CLmin), and free-molecule receptors in the cytoplasm, such as cGAS, IFI16, STING, DAI, and soon. In the present study, the innate immune response against SARS-CoV-2 is defined assuming that an optimal or adequate innate immune response eliminates in its entirety (100%) the pathogens with which it interacts^13^, and a suboptimal IIR is characterized by fail to control the virus in some degree. Under these conditions, IIR is defined as a characteristic function from [0,1] to [0,1], as follows:

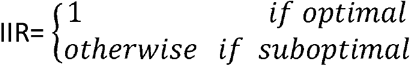

Now, for suboptimal Innate immune response (IIR<1), three categories (quartiles) are defined inspired by the latest clinical reports for COVID-19^6^,

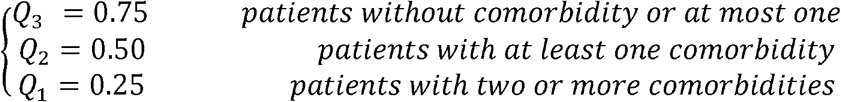

Meaning respectively, up to 75% efficiency in the clearing of the pathogen, up to 50% and up to 25%. On the other hand, the initial Viral load (α) and the viral burden (α), are expressed, respectively, in log_10_ copies/ml*hour, in short (copies/ml*h), and log_10_ copies/ml, in short (copies/ml) and following To KK-W et. al.^14^, their values are chosen in the real interval [1-10]. The interaction rate (*β* – 0.96) between the virus particles and the immune effector cells *(vi)* is obtained from Hernandez-Vargas & Velasco-Hernandez^15^, who derived it by fitting numerical estimations comparing the performance of their mathematical model with the data obtained from nine patients with COVID-19.

The innate immune system inhibits virus replication, promotes virus clearance, induces tissue repair, and triggers a prolonged adaptive immune response against the viruses, the latter is represented in the present model by the population of immune effector cells (i).

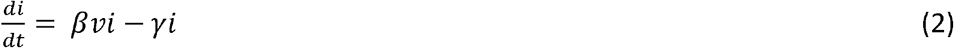

Equation (2) represent the dynamic of immune effector cells as a function of the interaction rate *β* between the virus particles with the immune effector cells (*vi*), and the death rate (*γ*) of immune effector cells. Regarding the immune effector cells (i), their values are expressed as *log*_10_ of the number of effector cells /ml. The death rate of immune effector cells (*γ* = 0.11) is taken from Macallan D et.al.^16^, who measured, using In vivo labeling with^2^H_2_-Glucose and cell sorting methods, the proliferation and disappearance of immune effector cells in healthy human volunteers.

### Cytokine release

Cytokines play a central role in limiting the viral spreading within the host during the early phases of the disease. The dynamic of cytokines (c) release is represented by equation (3).

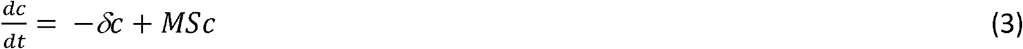

Assuming that the dynamic behavior of cytokine release, follows a mass action law which depends on the product of the stimulus S’ by the production rate *M* of cytokines by these cells. *S* is defined as the effect of immune cells activity on cytokine production, and *M*, following, Waito et.al.^17^, is defined as the maximum cytokine production rate expressed as a percentage. On the other hand, in the absence of these stimulated producers, the number of cytokines will rapidly decline at a rate *δ,* defined in function of the effect of intensity of virus exposure on cytokine concentration, assuming that viral decay exerts an appreciable influence on cytokine decline^18^. Finally, to record the dynamic of cytokine release the Log_10_ of average cytokine concentration is used.

### Systems dynamics approach

To develop a versatile strategy to study the complex issues of the within-host SARS-CoV-2 kinetic model, a system dynamic approach^19,20^ is built through the identification and description of the different parameters, components, feedbacks and processes that drive system behavior **(Table 1**). The Stock Flow diagram, corresponding to the differential equations of the model, is presented in **Appendix 1.**

**Table 1.**
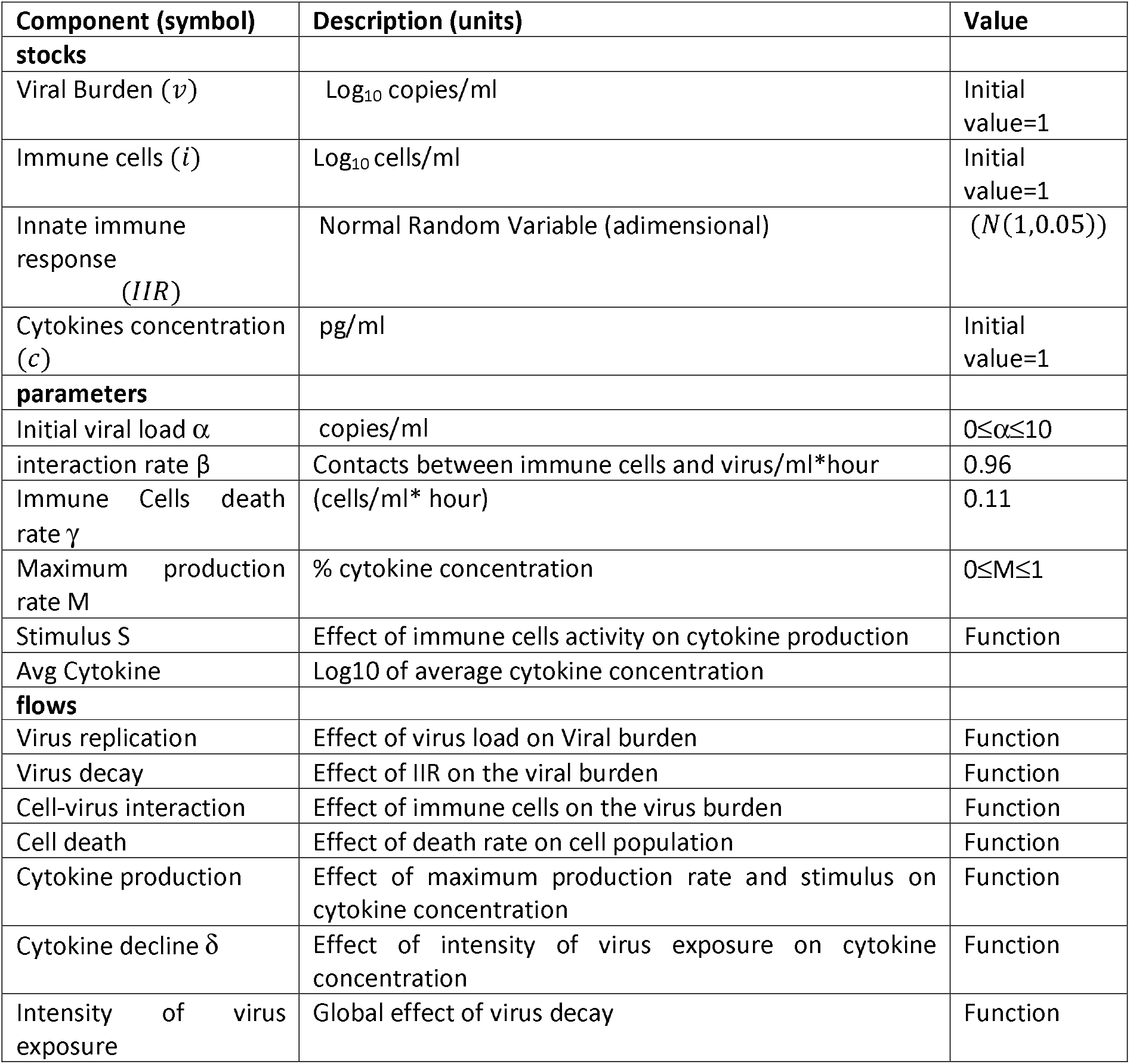
System dynamic model parameters

## Results

The dynamic simulation of the differential equation model (1), (2) and (3), under the conditions and parameters specified in **Table 1,** which seeks to represent the interactions between SARS-CoV-2 and the immune cells, gives rise to a series of interesting results. The first of them is related to the conditions that determine the occurrence or not of a periodic behavior associated with the dynamics in time of the immune response effector cells and that of the viral particle population, similar to the behavior of a predator-prey system (see **figure 1**).

**Legend Figure 1.**
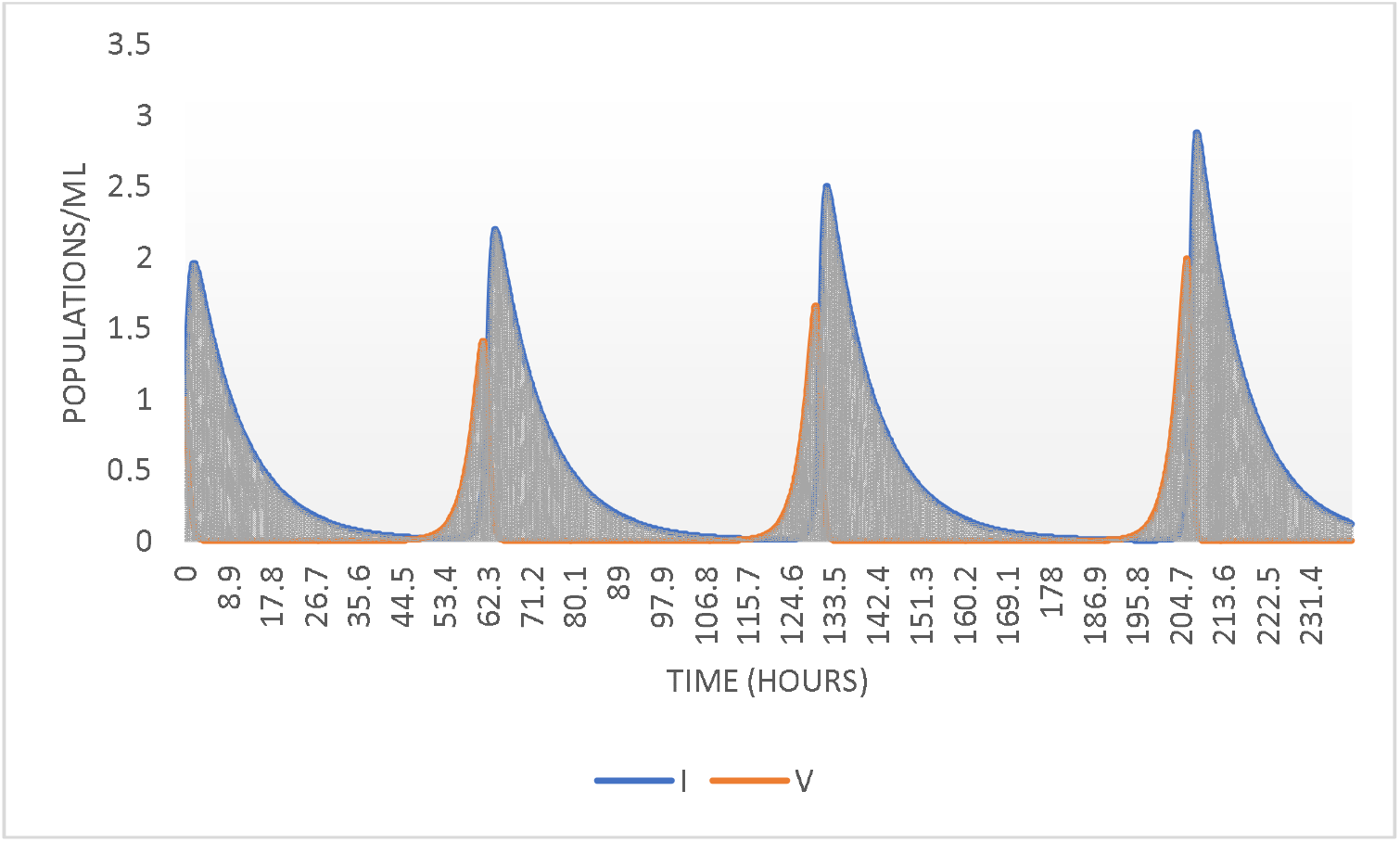
The coupled dynamics of immune cells and virus burden resembling recurrence of positive of SARS-CoV-2 RNA in COVID-19 patients.

As can be seen in **Table 2,** in cases of innate deficient or suboptimal immune response^21^, i.e. 0.1≤IIR<1, there is a phenomenon of viral shedding following an increasing oscillatory behavior, both in the frequency and level of viral load (copies/ml), as the IIR is lower and the initial viral load α is between 1 and 0.5 copies/ml. It is interesting to note that, with lower viremia, for the present case, 0.1 ≤ *α* ≤0.5 copies/ml, virus shedding events occur only when IIR= 0.1. The above provides an explanation for clinical observation related to positive RT-PCR results in patients who have recovered from COVID-19^22,23,24^. Following Li and collaborators^24^, the prolonged shedding of virus from recovered patients is obviously not an isolated phenomenon, but an integral component of the interaction dynamics between SARS-CoV-2 and the immune cells.

**Table 2.**
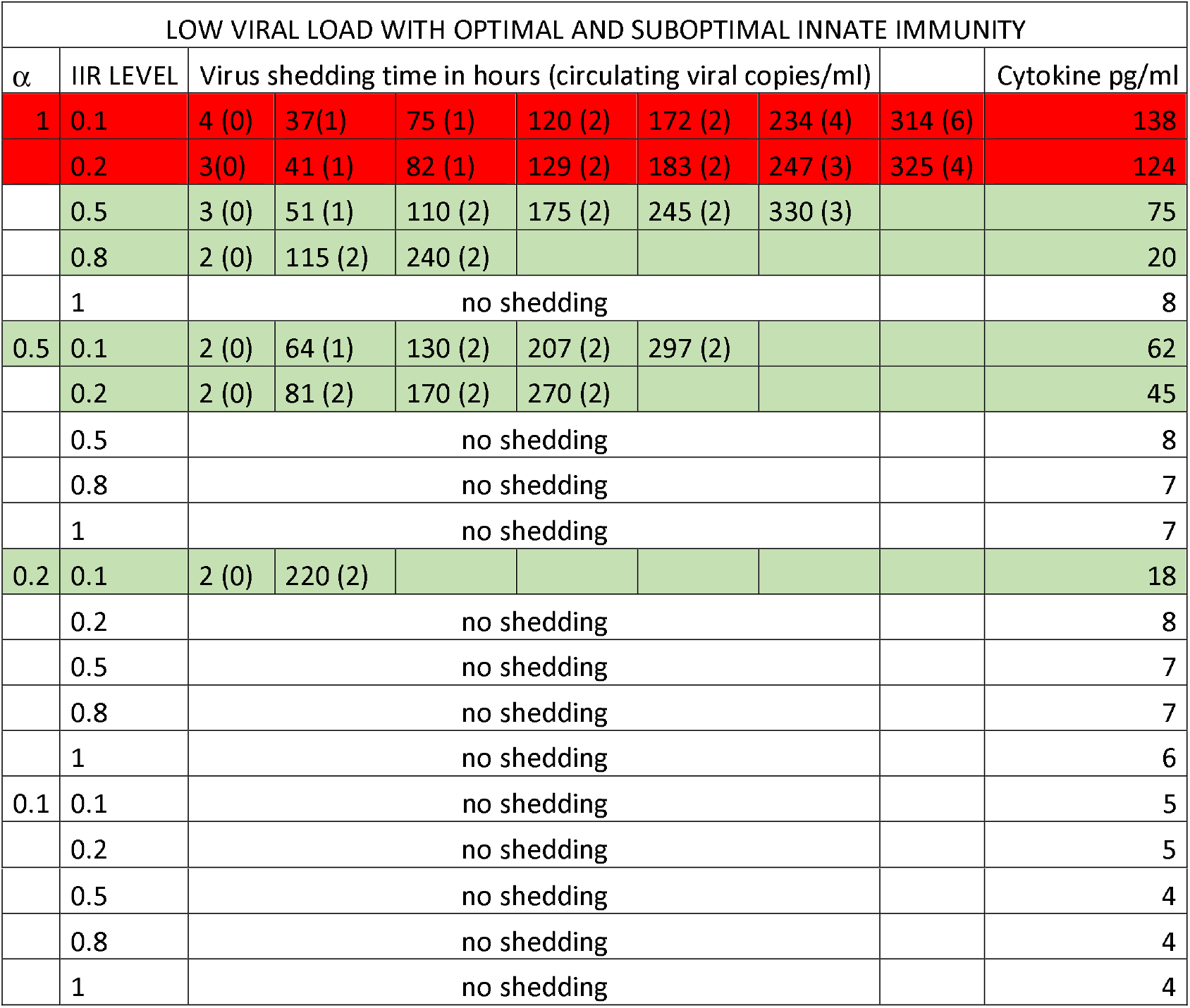
Scenario with low viral load and optimal and suboptimal innate immunity. The acute or severe symptomatic individuals corresponds with those lines in red. Asymptomatic or mild asymptomatic individuals are indicated with green lines. The white lines correspond to patients who get infected but do not get the disease.

The numerical results seem to indicate that, with low viral loads, even if the individual does not have an optimal IIR, it is possible not to develop COVID-19 disease, a fact that evidently reinforces the benefits of the social distancing policies and hygiene measures associated with the use of mouthpieces, among others, that have been adopted throughout the world.

It is also observed in **Table 2** that if IIR=1, the infection does not progress from any viral load equal to or less than 1. However, when *a* is greater than 1, it is found **(Table 3),** that even with IIR=1, i.e. optimal innate immunity, it is possible to generate an oscillatory shedding dynamic of infection, at least for *2≤a* ≤4 copies/ml. This would partly explain the high rate of infection in healthy hospital staff, because they are on the front line of the battle against the coronavirus, usually in an environment overexposed to the pathogen. Interestingly, for 4≤ *α* ≤10 copies/ml, there is no viral recurrence, which would indicate that, when the initial infection is carried out by high viral loads, some mitigation event occur, which also seems to happen even under conditions of low capacity of innate immune response (IIR=0.1) as shown in **Table 4.**

**Table 3.**
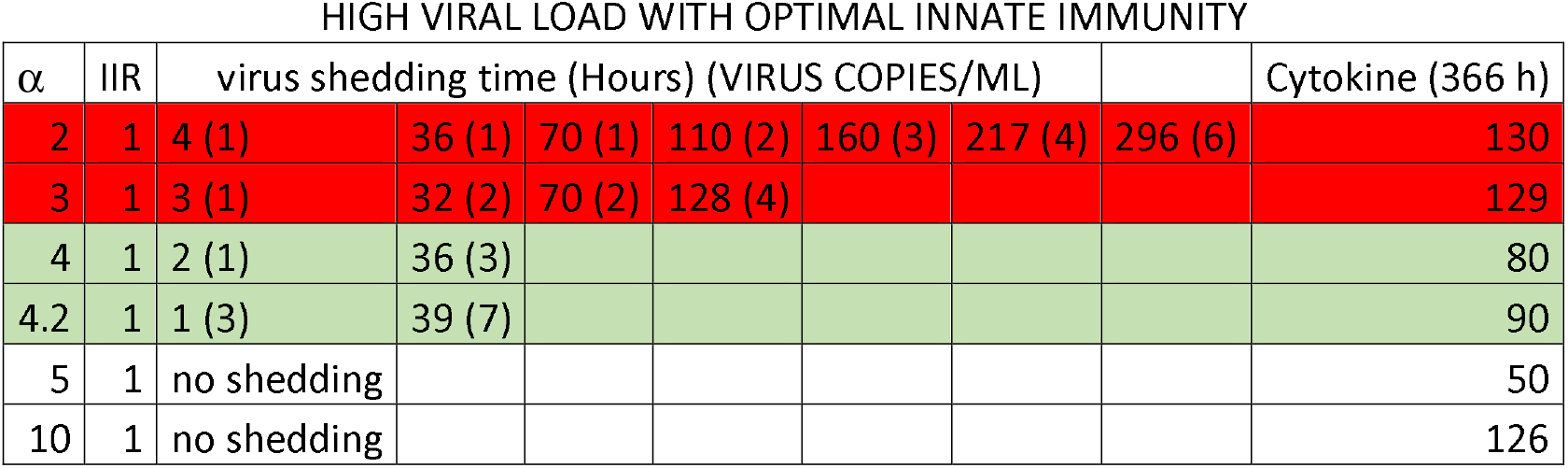
Scenario with high viral load with optimal innate immunity. When infection is contracted from high viral doses, it is possible to become ill with COVID-19, even if they have an adequate innate immune response, as highlighted in red. Individuals highlighted in green could corresponds to super-spreaders.

**Table 4.** Scenario with high viral load with a deficient innate immunity. The worst scenario, high viral load with deficient IIR, corresponding probably with severe symptomatic individuals. Note the high levels of cytokines in these cases. Interestingly, with α>4 copies/ml, a counterintuitive result appears, individuals who receive a high initial viral load, but do not develop the disease.

| HIGH VIRAL LOAD WITH DEFICIENT INNATE IMMUNITY |  |  |  |  |  |  |  |
| --- | --- | --- | --- | --- | --- | --- | --- |
| $\alpha$ | IIR | virus shedding time (hours) (VIRUS COPIES/ML) | | | | | Cytokine (366 h) |
| 2 | 0.1 | 3 (0) | 30 (2) | 67 (3) | 117 (5) | 207 (11) | 198 |
| 4 | 0.1 | 1 (4) | 53 (14) |  |  |  | 145 |
| 5 | 0.1 | no shedding |  |  |  |  | 66 |
| 10 | 0.1 | no shedding |  |  |  |  | 166 |

On the other hand, when the dynamics of cytokine levels are taken into consideration, the first thing that should be emphasized is that, under infection conditions characterized by the occurrence of oscillatory virus shedding, the concentration of cytokines has a continuous increasing behavior as can be seen in **Figure 2;** thus the cytokine storm can also be described as a cytokine cascade. Furthermore, the model suggests that the cytokine storm syndrome is also a result of a positive feedback between the two populations, under the specific combinations of initial viral load numbers and innate immune system status, since i) as can be seen in **Table 2,** with *a* =1 and 0.1≤IIR≤0.2, at the end of the simulation period of 336 hours, cytokine concentrations are high, between 124 and 138 pg/ml ii) in **table 3,** when *1≤a* ≤4 copies/ml, a final concentration of 130 pg/ml is reached, even when individuals have an optimal innate immune response. Finally, in **Table 4,** the worst-case scenario is observed, with a high initial viral load and a poor innate immune response, where at the end of 336 hours, a concentration of cytokines of 194 pg/ml is reached. It is possible to propose the hypothesis that individuals who meet the conditions described above of viral load, IIR status and cytokine concentration, could correspond to severe or critical symptomatic patients.

**Legend Figure 2.**
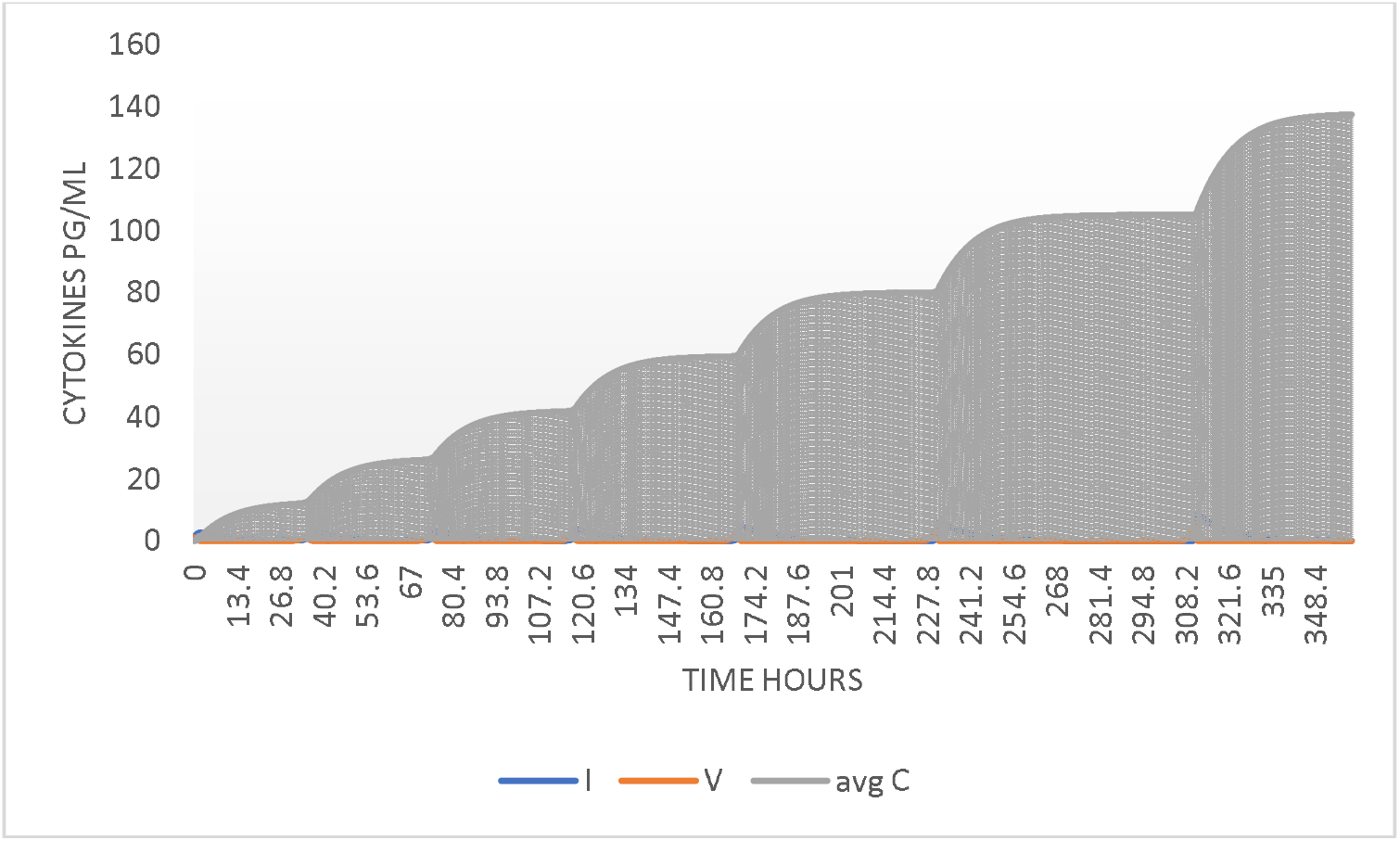
Increasing cascade of cytokines in patients with deficient innate immune response IIR<1 with 1 ≤ *α* ≤ 4. Some of these patients surely die, because the pro-inflammatory cytokines are detrimental.

In the literature on the clinical course of severe and critical COVID-19 patients^26^, the deregulated production of the pro-inflammatory cytokines contributes to inflammation, tissue damages and in acute COVID-19 patients, deposition of fibrin and platelet microthrombi in the lung vasculature, which contribute to progressive respiratory dysfunction and eventually to death. It’s the fine tuning between the molecules restricting the growth of pathogens and pro-inflammatory molecules which maintains the balance and helps in running the host system effectively, which seems to depend on the initial conditions of viral load and innate immune status.

The contribution of asymptomatic persons with MERS-CoV or SARS-CoV-2 to the transmission is not well characterized^27^. Those asymptomatic cases may play a role in the transmission and thus pose a significant infection control challenge^28^. Taking into account the results of the model related to cytokine concentrations at the end of 366 hours, and the data from the study of cytokine profiles in patients infected by SARS-CoV-2^29^, it is possible to propose that individuals with cytokine levels below 90 pg/ml with low viral burden dynamics (*v*≤3 copies/ml) and IIR<0.8, correspond to mild asymptomatic patients (see **Table 2).** Finally, the model suggests a definition for super-spreaders such as those **(Table 3),** whose circulating viral burden *v* of 3 copies/ml or more, but cytokine concentration below 90 pg/ml.

The present work shed some light on the definition of asymptomatic and super-spreaders status regarding SARS-CoV-2 infection in humans, but their role in the transmission deserve further studies to examine the clinical course of those individuals, viral dynamics, viral loads and immune status, to estimate their real contribution to the transmission dynamic of SARS-CoV-2. Such studies will enhance the understanding of the pathogenesis of these emerging viruses and will inform policy makers to make scientifically sound recommendations.

### Limitations

Some limitations of the study are the following. First, it is assumed that the interaction between viral particles and immune response effecting cells is of a symmetrical nature, when in fact it is mediated by the invasion of the virus into the epithelial cells of the lungs, which generates the stimulus that causes their interaction with the immune cells. On the other hand, it is assumed that the innate immune response acts as a simple characteristic function, which oversimplifies the chain of events related to the non-specific response, but allows the obtaining of a set of interesting results that explain some clinical observations associated with COVI-19, such as the cytokine storm.

## Conclusions

The present study hypothesizes over the conditions that characterize the fraction of the population which get infected by SARS-CoV2; the asymptomatic patients as those with optimal Innate immune response, exposed at low virus loads and controlling the viremia few hours after infection. These could be not infective when maintain contact with other people. The asymptomatic or symptomatic mildly infected patients, as those possessing a low viral burden with shedding episodes, given their suboptimal innate immune response and, with a cytokine level between 18 and 80 pg/ml. these patients could be infective to other people. In this category could be classified those patients, that although possesses an adequate innate immune response, were exposed to high virus loads and present virus shedding, but maintains cytokines level below 50 pg/ml. these individuals could corresponds to a super-spreaders group. Finally, the set of acute or severe symptomatic patients, characterized by deficient innate immune response, suffering viral shedding episodes frequently and with an increasing concentration of cytokines of 130 pg/ml or more. Some of these patients surely die, because the pro-inflammatory cytokines are detrimental, causing the cytokine storm with increasing viral load, tissue damages and pathophysiology.

## Ethics statements

No animal studies are presented in this manuscript.

No human studies are presented in this manuscript

No potentially identifiable human images or data is presented in this study.

## Data availability statement

All datasets presented in this study are included in the article/ supplementary files

---

**ANNEX 1.** STOCK FLOW DIAGRAM OF WITHIN-HOTS DYNAMIC OF WITHIN-HOST VIRAL KINETICS OF CORONAVIRUS (SARS CoV-2) IN HUMANS.

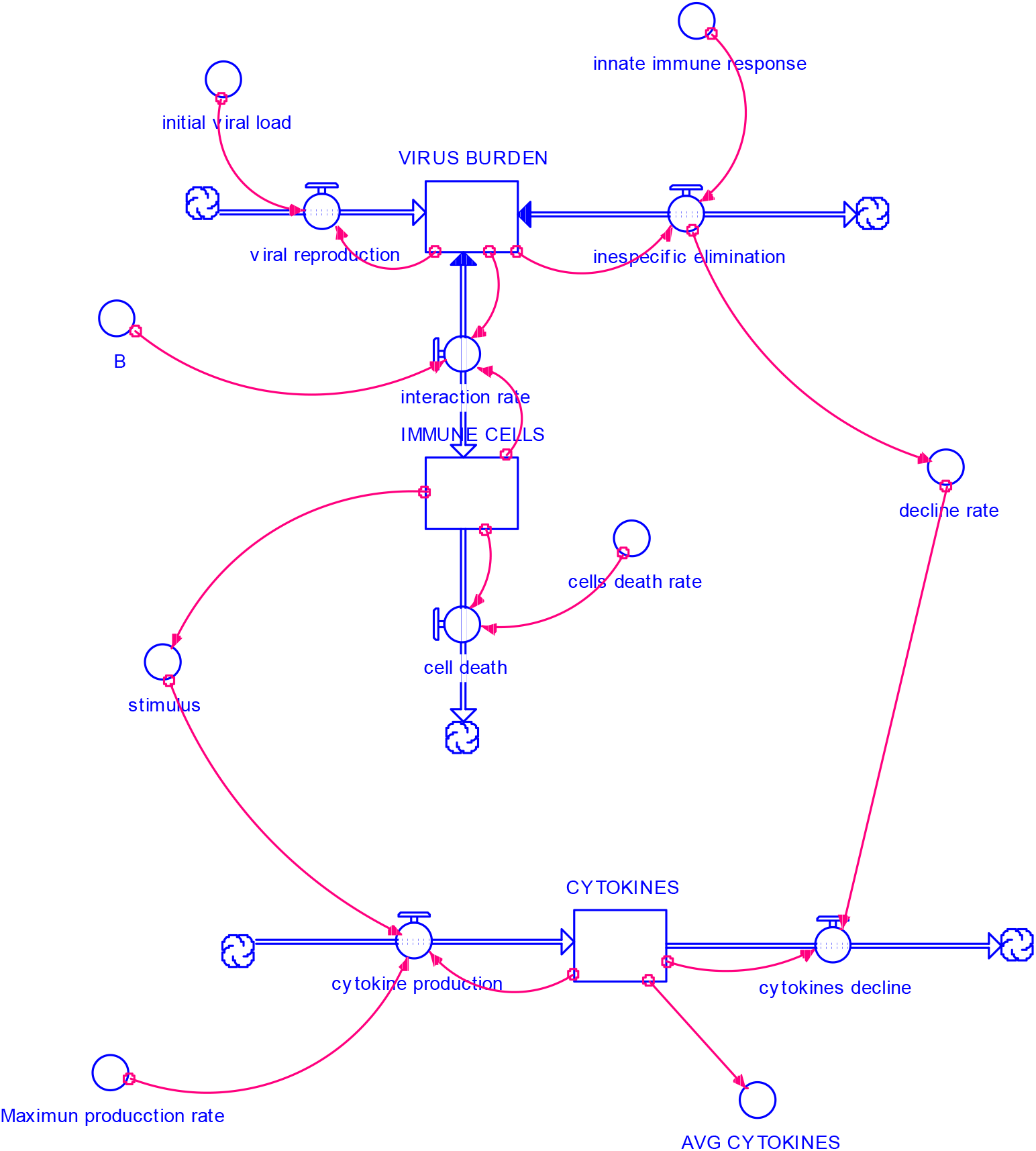

## References

1) Li, Q., Guan, X., P. Wu, P., et.al. Early Transmission Dynamics in Wuhan, China, of Novel Coronavirus-Infected Pneumonia. N. Engl. J. Med. 2020. 382, 1199–1207. doi:10.1056/NEJMoa2001316pmid:31995857

2) World Health Organization. Novel Coronavirus (2019-nCoV) situation report-2 [published online ahead of print January 21, 2020]. https://www.who.int/docs/default-source/coronaviruse/situationreports/20200122-sitrep-2-2019-ncov.pdf

3) Giordano, G., Blanchini, F., Bruno, R., Colaneri, P., Di Filippo, A., Angela Di Matteo, A. Colaner, M. Modelling the COVID-19 epidemic and implementation of population-wide interventions in Italy. 2020. Nature Medicine, https://doi.org/10.1038/s41591020-0883-7.

4) Wadman, M., Couzin-Frankel, J., Kaiser, J., Matacic, C. How does coronavirus kill? Clinicians trace a ferocious rampage through the body, from brain to toes. 2020. Science. doi:10.1126/science.abc3208.

5) Netea, M., Quintin J. and van der Meer, J. Trained immunity: a memory for innate host defense. 2011. Cell Host Microbe 9, 355–361.

6) Chen, N. et al. Epidemiological and Clinical Characteristics of 99 Cases of 2019 Novel Coronavirus Pneumonia in Wuhan, China: A Descriptive Study. 2020. The Lancet, vol. 395, 10223, 507–513., doi:10.1016/s01406736(20)30211-7.

7) Liang, K. Mathematical model of infection kinetics and its analysis for COVID-19, SARS and MERS. 2020. Infection genetics and Evolution. https://doi.org/10.1016/j.meegid.2020.104306

8) Lia, C., Xub, J., Liuc, J., and Zhou, Y., The within-host viral kinetics of SARS-CoV-2. 2020. bioRxiv preprint doi. https://doi.org/10.1101/2020.02.29.965418doi

9) Li, G., Fan Y., Lai Y., et.al. Coronavirus infections and immune responses. J. Med. Virol. 2020; 1–9. DOI: 10.1002/jmv.25685.

10) Lauer, S., Kyra, H., Grantz, B., Qifang, B. The Incubation Period of Coronavirus Disease 2019 (COVID-19) From Publicly Reported Confirmed Cases: Estimation and Application. 2020 Ann Intern Med. 172(9):577–582.

11) Akira S, Uematsu S, Takeuchi O. Pathogen recognition and innate immunity. Cell. 2006; 124(4):783–80.

12) Frieman, M., Mark Heiseb, M., and Bari, R. SARS coronavirus and innate immunity. 2008. Virus Research 133, 101–112.

13) Netea, MG., and Josten, LAB. Trained Immunity and Local Innate Immune Memory in the Lung. 2018. Cell. 175, 6, 1463–1465.

14) To, KK., et.al. Temporal profiles of viral load in posterior oropharyngeal saliva samples and serum antibody responses during infection by SARS-CoV-2: an observational cohort study. 2020. Lancet Infect Dis. 20: 565–74

15) Hernandez-Vargas & Velasco-Hernandez. 2020, In-host Modelling of COVID-19 Kinetics in Humans. MedRxiv preprints, https://doi.org/10.1101/2020.03.26.20044487.

16) Macallan, D. et.al. Rapid turnover of effector memory CD4 T cells in healthy humans. 2004. J. Exp. Med. 200 (2), 255–260.

17) Waito, M., Walsh, SR., Rasiuk, A., Bridle, BW., and Willms, A. A Mathematical Model of Cytokine Dynamics During a Cytokine Storm. 2016. In J. Bélair et al. (eds.), Mathematical and Computational Approaches in Advancing Modern Science and Engineering, DOI 10.1007/978-3-319-30379-6_31. Springer International Publishing Switzerland.

18) McDade, T., Georgiev, A., and Kuzawa, C. Trade-offs between acquired and innate immune defenses in humans. 2016. Evolution, Medicine, and Public Health pp. 1–16 doi:10.1093/emph/eov033.

19) Bala, B., Arshad, M., Noh, K. System Dynamics Modelling and Simulation. 2017. Springer. DOI 10.1007/978-981-10-2045-2.

20) Ellner, S. and Gluckenheimer, J. 2006. Dynamic Models in Biology. Pinceton University Press. 329 pp.

21) Nelemans T., Kikkert, M. Viral innate immune evasion and the pathogenesis of emerging RNA virus infections. 2019. Viruses. 11(10):961.

22) Lan, L. et.al. positive RT-PCR tests results in patients recovered from COVID-19. 2020. JAMA. 323, 15, 1502–1503.

23) Yang, J. et.al. persistent viral RNA during recovery period of a patient with SARS-CoV-2 infection. 2020. J. Med. Virol. DOI: 10.1002/jmv.25940.

24) Li, N. et.al. prolonged SARS-CoV-19 RNA shedding: not a rare phenomenon. 2020. J. Med. Virol. DOI: 10.1002/jmv.25952.

25) Yang Liu, Y., Yan, L-M., Wan, L., Xiang, T-X., Le, A., Liu, J-M., Peiris, M., Poon, L., Zhang, W. Viral dynamics in mild and severe cases of COVID-19. 2020. Lancet Infect Dis. https://doi.org/10.1016/S1473-3099(20)30232-2.

26) More, J., and June, C. Cytokine release syndrome in severe COVID-19. 2020. Science 368, (6490), 473.

27) Al-Tawfiq, J. Asymptomatic corona virus infection: MERS-CoVandSARS-CoV-2. 2020. Travel Medicine and Infectious Disease, https://doi.org/10.1016/j.tmaid.2020.101608.

28) Rothe, C., Schunk, M., Sothmann, P., et.al. Transmission of 2019-nCoV Infection from an Asymptomatic Contact in Germany. 2020. N. Engl. J. Med. 382, 970–971. doi:10.1056/NEJMc2001468pmid:32003551.

29) Liu J, Li S, Liu J, et al. Longitudinal characteristics of lymphocyte responses and cytokine profiles in the peripheral blood of SARS-CoV-2 infected patients [published online ahead of print, 2020 Apr 18]. EBioMedicine. 2020; 102763. DOI: 10.1016/j.ebiom.2020.102763.

